# Phototaxis is a state-dependent behavioral sequence in *Hydra vulgaris*

**DOI:** 10.1101/2023.05.12.540432

**Authors:** Soonyoung Kim, Jacob T. Robinson

## Abstract

Understanding how internal states like satiety are connected to animal behavior is a fundamental question in neuroscience. *Hydra vulgaris*, a freshwater cnidarian with only eleven neuronal cell types, serves as a tractable model system for studying state-dependent behaviors. We find that starved *Hydra* consistently moves toward light, while fed *Hydra* do not. By modeling this behavior as a set of three sequences - head orientation, jump distance, and jump rate -we demonstrate that the satiety state only affects the rate of the animal jumping to a new position, while the orientation and jump distance are unaffected. These findings yield insights into how internal states in a simple organism, *Hydra*, affect specific elements of a behavior, and offer general principles for studying the relationship between state-dependent behaviors and their underlying molecular mechanisms.

## Introduction

Animal behavior is a dynamic process influenced by environmental cues and internal states [1]–[3]. In neuroscience and ethology, uncovering the mechanisms by which complex behaviors are regulated is essential and can provide clues to the mechanics of decision-making. Computational modeling in particular has been useful in identifying the components of behavioral patterns as well as investigating the underlying biomolecular and neural driving factors [4]–[6]. However, the challenge in dissecting complex state-dependent behaviors to establish mechanistic models inevitably increases with the complexity of the anatomy, the connectome and the breakdown of observable behavioral motifs.

*Hydra vulgaris*, a freshwater invertebrate, is a particularly tractable model organism for studying the mechanisms behind state-dependent behaviors thanks to their basic body plan, simple nerve net, and easily-identifiable behavioral repertoire. *Hydra*’s body has two layers of tissue, the ectoderm and endoderm, separated by an extracellular matrix with no apparent canonical organs [7]. It exhibits sensorimotor reflexes to external stimuli such as temperature change [8]–[12], mechanical touch [11], [13]–[15], chemicals [16]–[19] and light [14], [20]–[22]. Its radially-symmetric nervous system consists of only a few thousand neurons in the form of a nerve net distributed throughout the entire body, with only 11 constituent neuronal subtypes [23], [24]. Despite the diffuse neural architecture, there is evidence that neurons in the oral region (i.e., the head) are essential for information processing [13]. Another advantage that *Hydra* possesses is the distinct and isolated behavioral repertoire such as longitudinal and radial contraction, elongation, nodding, feeding, and somersaulting that can automatically be classified [25]. Moreover, functional neural circuits associated with specific behavioral motifs have been recently uncovered, providing opportunities to correlate behavior with neural activity [26]. Given *Hydra*’s position in the phylogenetic tree as a part of the sister group to bilaterians, studying the regulation of complex behaviors in *Hydra* can further develop our understanding of fundamental processes that have been conserved over millions of years.

One complex behavioral motif found in its natural state is somersaulting, *Hydra*’s main mode of locomotion, where it anchors itself with the foot, and sways around with their body column, head and tentacles until tumbling in the direction the head was pointing, moving their foot to a new location [27], [28]. An even more complex behavior is phototaxis, which is a goal directed behavior: moving toward or away from a light source. This behavior was first described by Trembley in 1744 [29]. It has since been documented in both the symbiotic green *Hydra* (e.g., *Hydra viridissima*) which bears photosynthetic algae and asymbiotic brown *Hydra* (e.g., *Hydra vulgaris*) [30], [31]. Feldman and Lenhoff suggested that phototaxis may be sensitive to food deprivation [32]. This was particularly intriguing because it implied that *Hydra* not only are capable of goal-directed behaviors, but also that they change their behavior depending on their internal state. However, these reports remain observational, lacking a rigorous quantitative and systematic analysis to reveal mechanistic explanations behind the potential state dependency.

Here we show the first quantitative analysis and behavioral model of satiety-dependent phototaxis in wild-type AEP Kiel *Hydra vulgaris*. To perform this study, we developed a low-cost, high-throughput imaging setup to record the animals’ behavior in the presence of a controlled light stimulus for an extended period of time. We discovered that *Hydra* consistently moves towards the light within 8 hours when starved (i.e., 14 days post feeding) but this phototaxis is attenuated when fed (i.e., 2 days post feeding). These data suggest that satiety is a key internal state regulating phototaxis. Having found this property, we then asked which aspect of *Hydra*’s phototactic behavior is specifically modulated by satiety. To answer this question we first extracted three behavioral elements of phototaxis, namely head orientation, jump distance, and jump rate. We found that satiety only changed the jump rate. We confirm that the jump rate is solely responsible for the observed differences in phototaxis using a mathematical model that simulates this behavior using a simple algorithm with the aforementioned biophysical parameters.

## Results

### Phototaxis is a robust and reproducible complex behavior in *Hydra vulgaris*

Although phototaxis had been reported anecdotally, we wanted to develop a method to quantify this behavior and confirm its reproducibility. One challenge is that *Hydra* moves incredibly slowly (∼0.8cm/hr). To account for their slow movements, we built a microfluidic setup that allows scalable and long-term imaging in a controlled environment (Fig. 1a,b, see Materials and Methods). To test for phototaxis, we exposed the *Hydra* placed in the microfluidic chamber to an optical gradient provided by a set of LEDs and an optical diffuser installed to one side of the chamber (Fig. 1a,c). We used DeepLabCut, a deep learning-based pose estimation algorithm [33], to obtain and track the coordinates of the body parts, namely foot, body column, and head (Fig. 1c). Due to *Hydra* mainly locomoting by somersaulting, detaching the foot and attaching it to a new location, we used the foot coordinates to determine the location of *Hydra* [27], [28].

**Figure 1.**
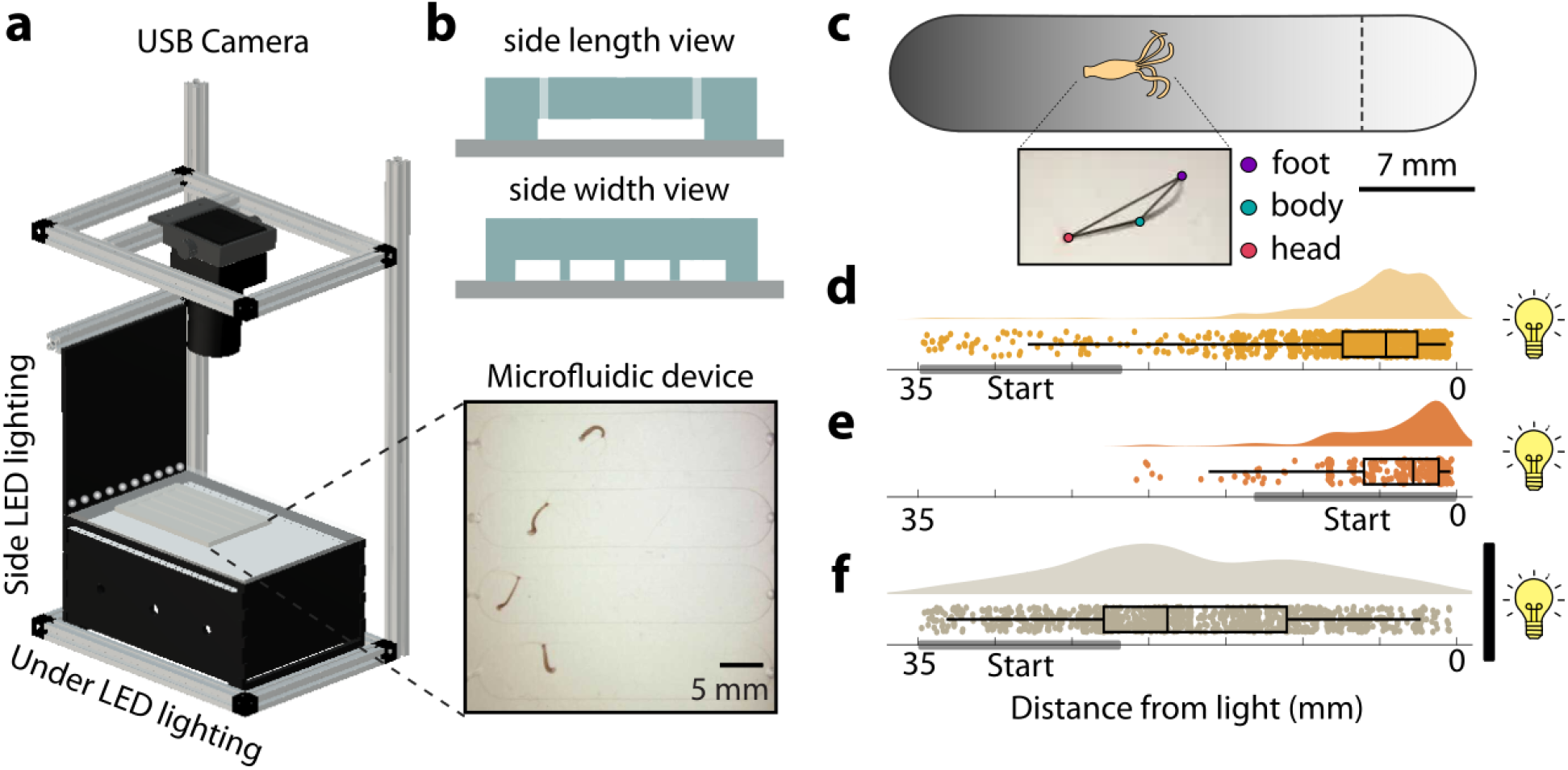
Phototaxis is a robust and reproducible behavior in *Hydra vulgaris*. (a) High-throughput, long-term imaging setup. Teensyduino-controlled LEDs are used for both side and under illumination. USB cameras are connected to a computer for recordings. Microfluidic devices containing Hydra are placed on the platform. (b) Microfluidic device overview. Top panel shows a typical schematic of the side length view, which depicts how the microfluidic lane is configured. The middle panel shows the width view of how lanes are distributed within one device. Bottom panel shows Hydra in each microfluidic lane. (c) Schematic of phototaxis experiment and labeling body points. A representative optical gradient across the microfluidic lanes is shown. The dashed line indicates the threshold for phototaxis. Inset depicts the labeled body points derived from DeepLabCut. Hourly foot location distribution of Hydra when (d) placed on the darker side of the chamber and exposed to light, pooled from N = 36, (e) placed on the brighter side of the chamber and exposed to light, pooled from N = 6, and (f) placed on the darker side of the chamber and light is blocked, pooled from N = 24. The gray boxes in (d)-(f) show the range of *Hydra*’s initial placement.

When initially placed on the darker side of the chamber, all 24 *Hydra* tested moved toward the light source (Fig. 1d). To verify that the phototaxis we observed was not an artifact of the animal’s preference to move to the opposite end from where they are placed, we also tested placing animals initially on the brighter side of the chamber (i.e., closest to the light source). In these experiments, *Hydra* stayed closer to the light source rather than move away (Fig. 1e). Given that *Hydra* responds to thermal stimulation, we also asked if the behavior could be driven by a thermal gradient generated by the LEDs rather than an optical gradient. We repeated the experiment using an optical diffuser painted black, which allowed the thermal diffusion to carry through while blocking the photon-emitting light. In this case, *Hydra* roamed around the chamber without any apparent direction (Fig. 1f), suggesting that the phototaxis behavior we observed is not an artifact of thermal differences of the geometry of the chamber and is in fact light-driven.

### Phototaxis in *Hydra* is satiety-dependent

In the course of maintaining *Hydra* we observed that when we had not fed the animals recently, they seemed to be more attracted to light. This unexpected result led us to ask if *Hydra* phototaxis is a state-dependent behavior. Considering that satiety level (i.e., hunger) is one of the major factors in driving behavioral shifts across the animal kingdom [34]–[38], we sought to investigate if this was also the case for phototaxis in *Hydra*. We divided an homogenous *Hydra vulgaris* AEP Kiel population into two groups, namely “*starved*” and “*fed*.” *Hydra* in the *starved* group were given any food for 14 days while the animals in the *fed* group were fed until 2 days prior to phototaxis recordings. For each group, *Hydra* were placed on the darker side of the chamber at the beginning of the experiment. We found that *Hydra*’s phototactic behavior was satiety dependent, with only the *starved Hydra* locomoting towards the light within the first 8 hours, and staying within the vicinity afterwards (Fig. 2a, Supplemental video S1). Fig. 2c shows representative traces of the foot locations over the course of the first 8 hours, where *Hydra* moves with strong directionality. In contrast, the phototactic behavior was drastically attenuated in the *fed* animals (Fig. 2b, Supplemental video S2), while some *Hydra* phototaxed, most either roamed with no apparent sense of direction with reduced movement, or barely locomoted at all (Fig. 2d).

**Figure 2.**
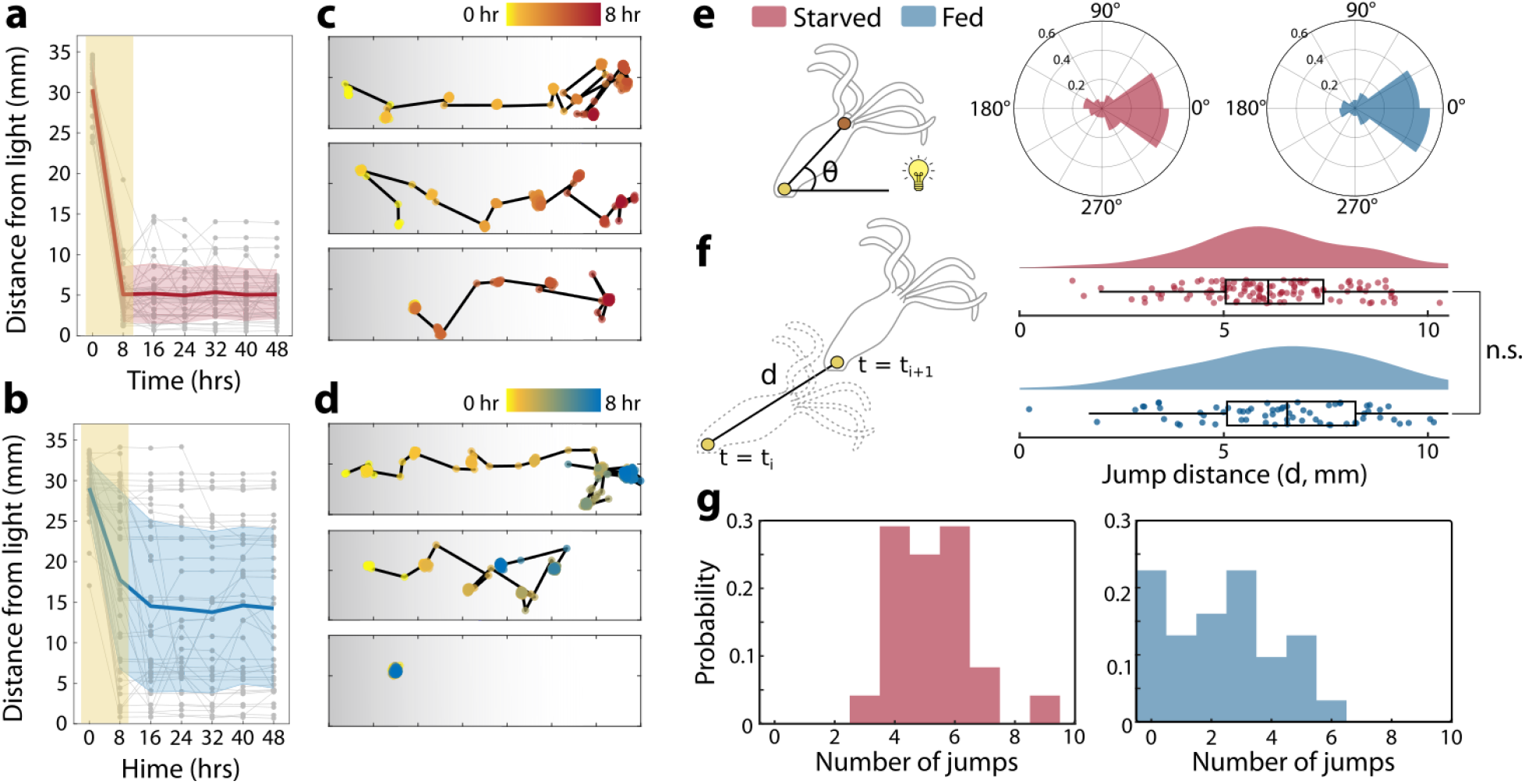
Satiety attenuates the phototactic behavior in *Hydra* with reduced jump rate. (a) Foot location of the *starved Hydra* over the course of 48 hours, sampled at 0, 8, 16, 24, 32, 40 and 48 hr time points (N = 24). (b) Foot location of the *fed Hydra* over the course of 48 hours, sampled at 0, 8, 16, 24, 32, 40 and 48 hr time points (N = 31). With *Hydra* initially placed on the darker side of the chamber, representative foot location traces from the first 8 hours of experiment of (c) *starved Hydra* and (d) *fed Hydra*. (e) Left panel shows how head orientation is calculated. Right panels show the head orientation distribution of *starved Hydra* (shown in pink) and *fed Hydra* (shown in blue). (f) Left panel shows how the jump distance is calculated. Right panels show the jump distance distribution of *starved Hydra* (shown in pink) and *fed Hydra* (shown in blue). (n.s. = not significant, Mann-Whitney U Test.) (g) Distribution of number of jumps of *starved Hydra* (shown in pink) and *fed Hydra* (shown in blue).

To understand the underlying mechanism behind the phototaxis, we focused our analysis on the first 8 hours of the experiments, which we found to be sufficient for *starved Hydra* to reach the light. We sought to identify key constituent behavioral parameters in phototaxis, and which of those were affected by the satiety level. *Hydra* mainly uses somersaulting to move to a new location, by tumbling in the direction its head was last pointing. It is also believed that photoreceptors or opsins are primarily located in the head and tentacles [39], [40], which is pertinent to light perception. Based on the prior knowledge, we extracted three parameters, namely head orientation, jump rate, and jump distance.

First, we observed the head orientation with respect to the light source to determine if the distinct behavioral patterns stem from a difference in light perception between *starved* and *fed Hydra*. Head orientation (θ) was calculated using the coordinates obtained from DeepLabCut (Fig. 1c), as the angle between a vector connecting the hypostome and the foot, and a vector connecting the hypostome and the light source (left panel in Fig. 2e). Interestingly, we found that the differences in collective head orientation for both *starved* and *fed Hydra* were indistinguishable (right panels of Fig. 2e), suggesting that the head orientation was not the direct factor in the behavioral difference. Next, we asked if the shift in the behavior arises from the difference in the jump distance. The jump distance was calculated as the distance between the location of the foot before and after translocation. We observed no statistically significant difference between the jump distance distributions (D) of *starved* and *fed* animals (Fig. 2f). In fact, we found that the difference lies in the number of jumps (Fig. 2g), which translates to jump rate. Representative foot traces of *starved* and *fed Hydra* (Fig. 2c,d) illustrate how *starved* animals generally display a higher number of jumps compared to *fed Hydra*. Roughly half of the *fed* animals translocated three times or less per trial.

### Phototaxis in *Hydra* can be synthetically created using a reduced set of parameters

Having identified the jump rate as the most critical of the three behavioral parameters involved in phototaxis, we considered if that parameter alone was sufficient to explain satiety-dependent phototaxis. To answer this question, we used the three aforementioned parameters and a simple algorithm (Supplemental Fig. 2) to generate synthetic phototaxis data sets. Each simulated *“Hydra*” was given a head orientation, a jump rate, and a jump distance, where the next foot location was determined by this sequence for every iteration until it crossed the phototaxis threshold or reached the termination time of 8 hours (Fig. 3a). Apart from the jump rate (λ), the parameters are the same between the synthetic *starved* and *fed* groups, and each synthetic *Hydra* was given a starting point between 25 and 35 mm from the light source.

**Figure 3.**
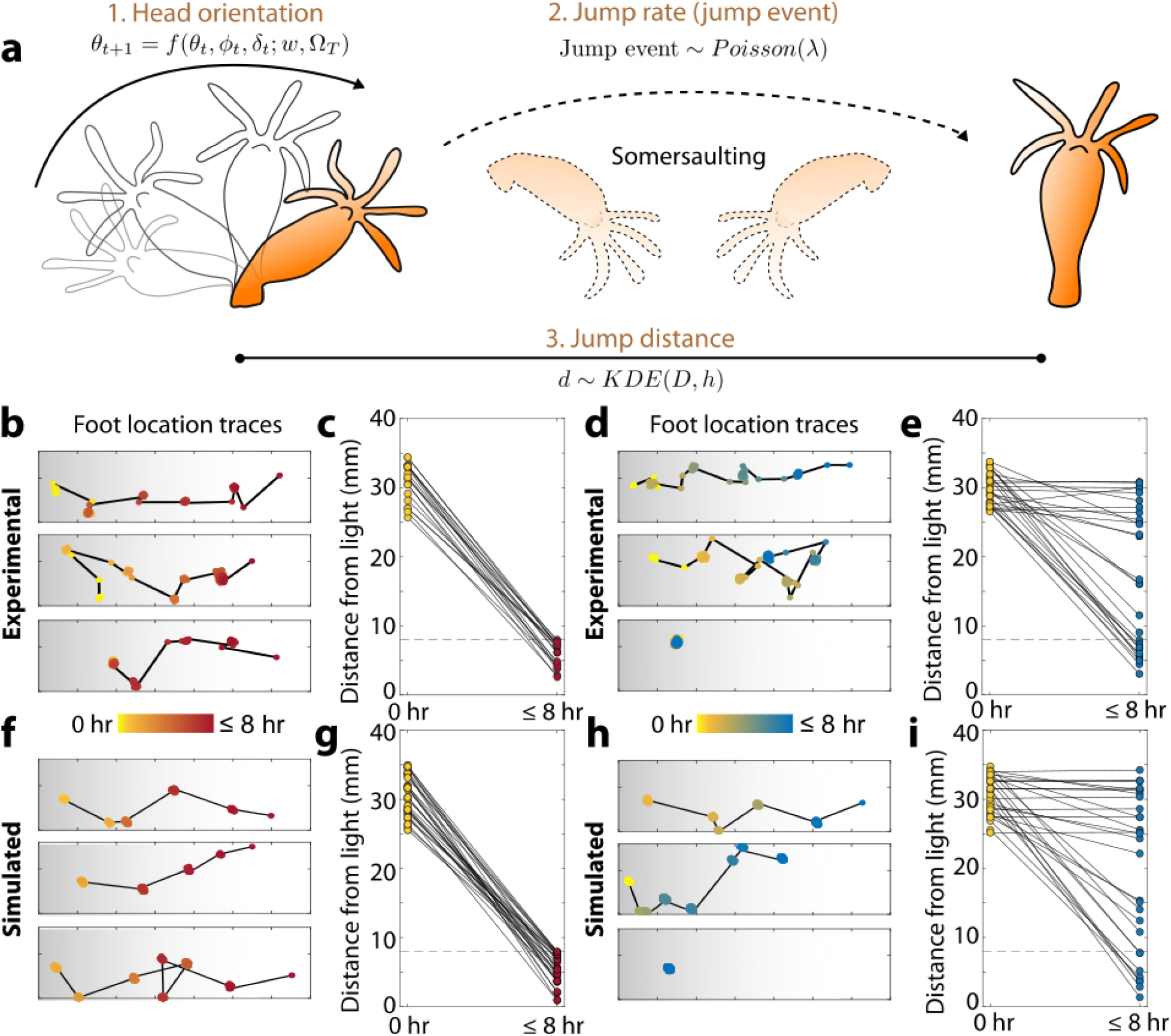
Modeling phototaxis in *Hydra* validates the jump probability is responsible for state dependency. (a) Schematic of the model using head orientation with biased correlated random walk (BCRW), Poisson process as the jumping decision-making, and jump distance distribution fitted from experimental data. Representative experimental foot location traces of (b) *starved Hydra* and (d) *fed Hydra*. Experimental foot location from start (0 hr) to termination (≤8 hrs) of (c) *starved Hydra* (N = 24) and (e) *fed Hydra* (N = 31). Simulated foot location traces of (f) *starved Hydra* and (h) *fed Hydra*. Representative simulated foot location from start (0 hr) to termination (≤8 hrs) of (g) *starved Hydra* (N = 30) and (i) *fed Hydra* (N = 30).

Since the head orientation (θ) smoothly progresses with changes in orientation (Δθ) sharply centered around 0° (Supplemental Fig. 1e), we adapted the concept of biased correlated random walk (BCRW) to model the head orientation. BCRW consists of two main terms: target that represents the bias (Ω_T_), and persistence that depicts correlation [41]–[43]. The target in this case is the light source, and persistence is the direction towards which *Hydra* was pointing in the immediate previous iteration. Bias was given the weight *w* and persistence (*1-w)*. Each term includes noise terms (φ and δ respectively) as well, for which we used the truncated location scale distribution fitted from the experimental data, yielding

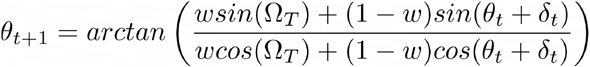

where t corresponds to a given iteration. After selecting the initial value for head orientation uniformly at random ranging from 0° to 360°, the progression followed the BCRW. The use of such a method resulted in similar head orientation trajectories between the experimental and simulated data for both *starved* and *fed* groups, indicating realistic modeling (Supplemental Fig. 1a-d). The jump rate was calculated from the experimental data, which differed depending on the starved or *fed* population. Due to the varying behavior in the *fed* animals (Fig. 2b,d), with roughly half of the *Hydra* executing a total of three jumps or less, we separately calculated jump rates for “Active” and “Inactive” *fed Hydra*. Given the nature of the somersaulting behavior where the time between jumps, or “events,” is random, we treated the phototactic movement as a Poisson process. Thus, the jump rate was calculated as λ, the Poisson variable, as follows:

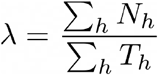

where N_h_ denotes the number of jumps for a given observed *Hydra*, and T_h_ denotes the total time for the same given *Hydra*. We also obtained the jump distance (d) from the distribution of the experimental data. Since the jump distance distributions for *starved* (D_s_) and *fed* (D_f_) *Hydra* are not statistically significantly different (Fig. 2f), the data points were pooled and fitted with a kernel density estimate using both data and bandwidth as inputs for sampling in simulations. When a jump event occurred, a jump distance was randomly sampled from the kernel-fitted pooled distribution D, and in the case of no jump, the jump distance was set to zero. Using these set of parameters, the next foot location was determined using the following equation:

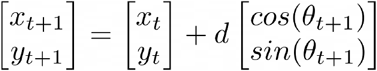

We simulated *Hydra’s* phototactic behavior with this minimal set of three variables and a simple algorithm. Quantitatively, the foot location traces between the experimental *starved Hydra* (Fig. 3b) and simulated *starved Hydra* (Fig. 3f) were comparable, with apparent biased movement towards the light. The simulated *Hydra* crossed the phototaxis threshold within the 8-hour limit (Fig. 3g), showing no differences with the experimental data (Fig. 3c). Similarly, for *fed Hydra* the foot location traces are analogous between simulated and experimental *fed Hydra* (Fig. 3d,h). The foot location distribution from start (0 hrs) to the termination of simulation (≤ 8 hrs) confirms similar patterns between the experimental (Fig. 3e) and simulated data set (Fig. 3i), with 2 subpopulations of Active and Inactive *fed Hydra*.

## Discussion

The ability to shift behaviors depending on internal states found in many species is an important feature for animal survival. Here, we demonstrate that despite being phylogenetically distant from most model organisms in neuroscience and having a basic nervous system with a rudimentary anatomy, asymbiotic *Hydra vulgaris* is capable of performing complex goal-directed behaviors that are modulated by internal states. Prior to this publication, phototaxis in brown *Hydra* such as *Hydra vulgaris* had only been described in anecdotal reports. Using a custom-designed imaging setup and microfluidic devices, we conclusively showed that *Hydra* indeed displays robust phototaxis. More importantly, the phototactic behavior was strengthened in the Starved population, whereas it was attenuated in the Fed population, which indicates that there are state-dependent internal processes that modulate neural circuit activity and behavior. By breaking down the phototactic behavioral motif into individual elements: head orientation, jump distance, and jump rate, we found that only the jump rate changed from *starved* to *fed Hydra*, and confirmed that this change alone is sufficient to explain our data when we simulated *Hydra* phototaxis in our model. This result provides a specific behavioral change that derives from satiety and shows where future work can focus to understand the molecular underpinnings of this state change.

We do not yet understand how *Hydra* senses where a light source is located. Prior reports have shown that *Hydra* can perceive and exhibit a sensorimotor reflex to light, and our results imply that *Hydra* can detect optical gradients without canonical visual organs, and thus possibly only with photoreceptors [32], [44]–[47]. One possibility for future work is to test if *Hydra* explores its environments by moving its photoreceptor-dense head and tentacles, storing the information over a short period of time, before somersaulting toward the strongest light stimuli. This is particularly interesting because it suggests that *Hydra* possesses memory-based decision-making capabilities, which has not been reported before. Future work should explore how *Hydra* may compute optical gradients, decide to move towards the light source based on this computation, and why *Hydra* stays at the brighter side of the chamber even after phototaxis is complete.

Another future direction is to investigate the biomolecular factors that drive satiety-dependent phototaxis in *Hydra* based on the demonstrated behavioral framework. *Hydra* genome has been fully sequenced [48], [49], and recent advances in molecular technology, especially single-cell RNA sequencing (scRNA-seq) provides a tremendous opportunity to probe the responsible genes and cell types that cause shifts in behavior [50], [51]. In conjunction with gene expression profiles, Assay for Transposase-Accessible Chromatin with high-throughput sequencing (ATAC-seq) can help reveal the epigenetic factors [49], [52]. Such findings can then further provide opportunities to behavioral modulation, perturbing the signaling and processing pathways of phototaxis to possibly disable or alter the phototactic predisposition. Together, these studies may help establish a comprehensive understanding of internal state-driven complex behaviors in a simple yet intricate organism.

With this future work it may be possible to understand why *Hydra*, particularly when starved, are attracted to light. This phenomenon is observed in many organisms, including bacteria and other invertebrates [53], [54]. Light perception is deemed necessary for survival, such as finding food, shelter and avoiding predators [55], [56]. The evidence that cnidocyte discharge in *Hydra* is mediated by optical input [21], coupled with the fact that *Hydra* captures its prey using their cnidocytes, suggests that phototaxis in *Hydra* may be a result of food search. Similar to *Drosophila* that has two distinct dopaminergic neural circuits that regulate foraging behaviors depending on the nutritional state [38], there is a possibility that *Hydra* may also have a circuit yet characterized that governs satiety-controlled behaviors.

The aspect of this work that is the most exciting to the authors is that *Hydra*, despite its relatively simple neuronal repertoire and architecture, is capable of goal-directed behaviors that are altered by internal states. The fact that this relatively sophisticated behavior is manifested in such a simple nervous system raises the prospect of developing a complete quantitative understanding of behavior that spans the molecular, cellular and organismal scales.

## Material and methods

### *Hydra vulgaris* strain and maintenance

All experiments were conducted on the *Hydra vulgaris* AEP Kiel strain. The animals were cultured using the protocol adapted from the laboratory of Robert Steele (University of California Irvine). *Hydra* were maintained at 18°C in an incubator with 12:12 hours of light:dark cycle controlled by a Teensyduino, and were fed three times a week with freshly-hatched *Artemia* nauplii (Brine Shrimp Direct). *Hydra* in the *starved* group were starved for 14 days, and those in the *fed* group were starved for two days prior to phototaxis experiments. *Hydra* were not reused unless explicitly stated otherwise.

### Microfluidic device fabrication and usage

All microfluidic chambers were fabricated using polydimethylsiloxane (PDMS) (Sylgard 184). The master mold, adapted from a previously designed lanes [19], were 3D-printed (Formlabs Form3) to have 6 lanes per device. After the inlet and outlet ports were punched with a 1.25 mm diameter biopsy, the molded PDMS was O_2_ plasma-bonded to a 75mm x 50mm glass slide (Corning 2947-75X50). After loading *Hydra* in the device following a previously-established protocol [8], [19], the top surface of the device including ports were covered with optically-clear adhesive well plate seals (Thermo Fisher Scientific AB1170) to minimize evaporation throughout the duration of the experiment. After each experiment, *Hydra* were unloaded from the device, and the devices were flushed with DI water and then soaked in 70% ethanol. After flushing the 70% ethanol out with DI water again, the devices were soaked in DI water and boiled at 150°C for ∼2 hours, and completely dried on hot plates until future use.

### High-throughput long-term imaging setup

The lighting setup used in the experiment was built using MakerBeams, custom-made PCB, LED strips (BTF-Lighting WS2812B), commercial USB camera (2.8-12mm varifocal with 8 megapixel), and custom-cut acrylic casing. The MakerBeams were used to build the skeleton, with a rectangular bottom and top structure connected by two vertical columns. One additional column and row was used for securing a side lighting panel. Acrylic was used to encase the under lighting panel and serve as a stage to place devices for imaging. An optical diffuser was attached to the stage and the side lighting panel for even diffusion. The LEDs were controlled with Teensyduino and TyCommander. The USB cameras were operated using the iSpy software. Each setup was housed in a cabinet with doors in order to prevent external light leakage.

### Phototaxis assay

Prior to the start of the experiment, each microfluidic device with *Hydra* inserted in them was placed on the imaging setup. For all experiments, one row of LEDs were turned on to maximum brightness (R:G:B 255:255:255) from the side lighting panel that yielded an optical gradient across the length of the microfluidic lanes. Under lighting at reduced brightness (R:G:B 1:1:1) without any gradient was additionally used for experiments where the light from the side panel needed to be blocked, in order to facilitate video recording. Timelapse videos were recorded at 1 frames per second for 48 hours. Foot coordinates (See Behavioral analysis and generating synthetic phototaxis dataset in Methods) were used to determine Hydra’s location at a given time.

### Behavioral analysis and generating synthetic phototaxis dataset

DeepLabCut [33] was used to quantify *Hydra*’s body coordinates. Locations annotated were basal disk (foot), center of the body column (body), and the hypostome (head). Mislabeled frames were manually re-labeled. The coordinates were extracted every 20 frames from the videos, to reduce the number of repeated frames due to *Hydra*’s slow movement. Custom MATLAB code was used for data analysis and modeling, including phototaxis simulations following the algorithm described in Results.

### Statistical analysis

Mann-Whitney U test (Wilcoxon rank sum test) was used to evaluate statistical significance between jump distances (Fig. 2f).

