## Supplemental Figure 1 for "Phototaxis is a state-dependent behavioral sequence in *Hydra vulgaris*"

### Supplemental Material

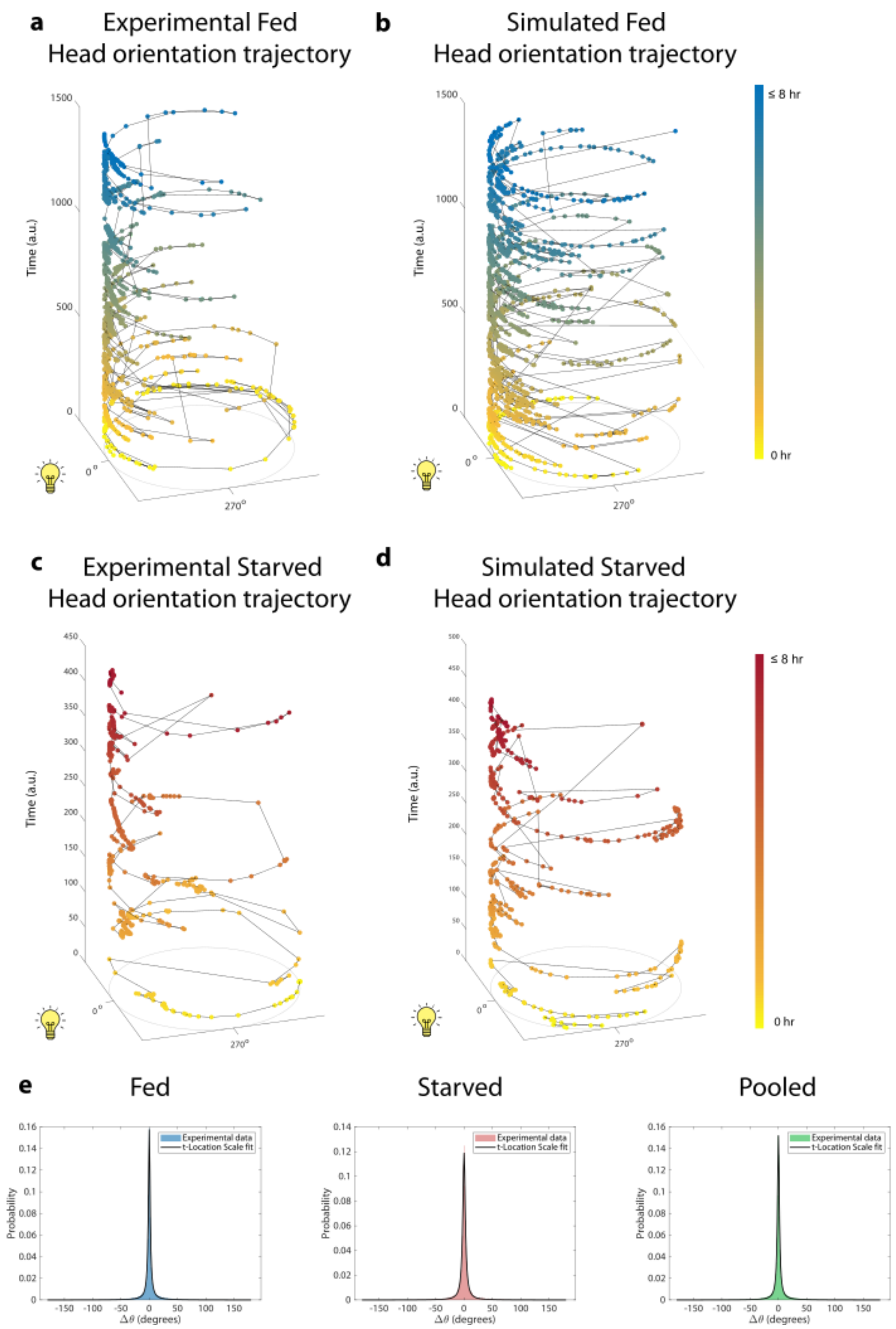


**Supplemental Figure 1. Head orientation trajectories.** (a) Head orientation trajectory of a representative experimental *fed Hydra*. (b) Head orientation trajectory of a representative synthetic *fed Hydra*. (c) Head orientation trajectory of a representative experimental *starved Hydra*. (d) Head orientation trajectory of a representative synthetic *starved Hydra*. (e) Differential head orientation for *fed* (left panel), *starved* (middle panel), and Pooled (right panel).

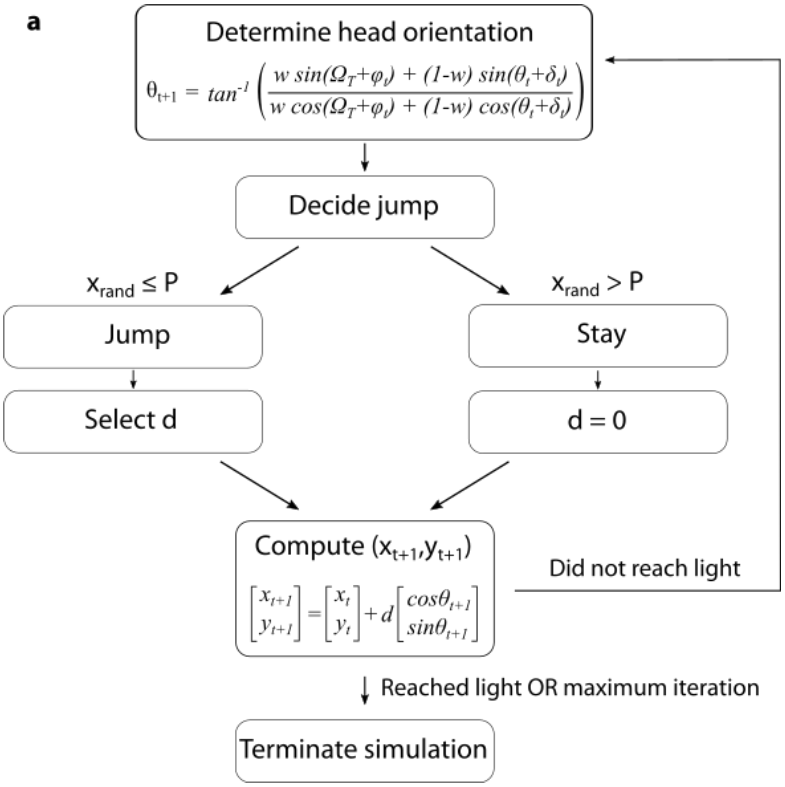


**Supplemental Figure 2. Flowchart describing the modeling process.**

**Supplemental Video S1. Representative 8-hour recording of *starved Hydra*.**

**Supplemental Video S2. Representative 8-hour recording of *fed Hydra*.**
